# Morphology and distribution of a new scale mechanoreceptor type in olive-headed sea snakes (*Hydrophis major*)

**DOI:** 10.1101/2025.01.28.635253

**Authors:** Alizée Wagner, Chad Johnson, Myoung Hoon Ha, Kate L. Sanders, Shaun P. Collin, Jenna M. Crowe-Riddell

## Abstract

Snakes are known for their superb sensory specialisations but are not widely appreciated for their sense of touch despite emerging evidence of tactile specialisation among sea snakes. This is partly due to the challenges in quantifying such small and numerous scale mechanoreceptors or ‘scale sensilla’ across individuals. By using a novel application of gel-based 3D profilometry (GelSight scanner) in combination with scanning electron-and light microscopy, we comprehensively quantified the morphology and distribution of scale mechanoreceptors in a sea snake, *Hydrophis major* (Hydrophiinae), for the first time. We discovered a new type of scale mechanoreceptor distinguished by its larger size (43.65 ± 22.57 μm height) and asymmetrical peak shape, created by a thickening in the cornified outer layers of the epidermis. Asymmetrical peaks contain a dermal papilla with central cells, indicative of meissner-like corpuscles that underlie smooth dome-shaped mechanoreceptors (31.88 ± 19.03 μm height) typically found in both sea snakes, but is positioned slightly off centre from the tallest point of the asymmetrical peak. Smooth domes are concentrated anterior-posteriorly on the head with the highest densities on the rostrum and nasal scales (9.83 ± 1.88 and 5.24 ± 2.85 per mm², respectively). Asymmetrical peaks are rarer; detected only on the dorsal and lateral sides of the head and most common on the postocular scale (mean density 1.26 ± 0.72 per mm²). We suggest functional differences in mechanosensory capabilities: asymmetrical peaks may serve primarily proprioceptive purposes whereas smooth domes are used for hydrodynamic reception.

## 1. INTRODUCTION

Tactile mechanoreception is a ubiquitous sense in animals. Although often considered a passive sense stimulated via direct contact with an object, many aquatically-foraging vertebrates use distal mechanoreceptors as an exploratory way to actively navigate within their environment and seek prey (Catania, 2012; Schneider et al., 2016). In mammals, for example, pinnipeds have highly vascularised whiskers (vibrissae) that they manipulate to detect the hydrodynamic trails generated by the fins of fish (Hanke et al., 2012). Specialised mechanoreceptors (Grandry corpuscles) on the bills of shore birds are used to probe for invertebrates in wet sand (Schneider et al., 2017), while non-avian reptiles, such as crocodilians have integumentary sense organs with underlying merkel cells that are concentrated along the jawline to detect ripples on the surface of the water caused by falling prey (Leitch & Catania, 2012; Soares, 2002).

Among snakes, tactile mechanoreception is partially mediated by small cutaneous organs called ‘scale sensilla’ that are distributed over the body but are concentrated in hundreds to thousands on the head (Jackson & Sharawy, 1980). At high magnification, these mechanoreceptors appear as raised ‘bumps’ over the scales and are created by clusters of innervated dermal cells called ‘dermal papillae’, which displace the overlying epidermis (Jackson et al., 1996; Von Düring & Miller, 1979). This underlying cellular structure is similar to that of Meissner corpuscles found in glabrous (hairless) skin in mammals. Scale mechanoreceptors are responsive to direct deformation of the skin via the displacement of these Meissner-like corpuscles (Jackson & Doetsch, 1977a; Proske, 1969). Tactile mechanoreception in terrestrial snakes is thought to be important for detecting the surrounding substrate during locomotion and for cloacal alignment during ‘tactile courtship’ (Jackson et al., 1996; Noble, 1934, 1937). Various aquatic snakes also use mechanoreception to forage for prey underwater, using modified scale structures that are assumed to detect water movement generated by prey (hydrodynamic reception).

Snakes are remarkably successful at living in aquatic habitats with multiple lineages independently evolving hunting strategies for capturing aquatic prey with the aid of touch (Murphy, 2012). File snakes (*Acrochordus* spp., Acrochordidae) are ambush predators that have small hair-like ‘scale sensillae’ likely used to detect fish movements in brackish/murky habitats (Povel & Van der Kooij, 1997). Similar to terrestrial snakes, these scale mechanoreceptors are distributed over the body but are more abundant on the head with up to seven per supralabial scale. Some scale mechanoreceptors are also more complex on the head, forming a series of branching structures with multiple fine hairs at their tip (Povel & Van der Kooij, 1997). Independently aquatic tentacled snakes (*Erpeton tentaculatum,* Homolopsidae) use paired and innervated ‘tentacles’ to detect the stereotyped escape S-response of fish as they attempt to evade the jaws of the snake (Catania, 2010; Catania et al., 2010; Winokur, 1977).

Sea snakes are a diverse radiation of front-fanged venomous snakes (∼70 species; Hydrophiinae), most of which spend their entire lives at sea and actively forage for fish (Crowe-Riddell et al., 2024). The sensory systems of sea snakes have adapted to the selection pressures of aquatic habitats including shifting the spectral sensitivity of their photoreceptor visual opsins to match the light environment at the depths they inhabit (Rossetto et al., 2024; Simões et al., 2020) and expanding the complement of chemoreceptor genes to detect odours that are encountered using tongue-flicking (Li et al., 2021). Scale mechanoreceptors in sea snakes are anatomically distinct from file snakes and tentacled snakes, and have been co-opted from terrestrial mechanoreceptors relatively recently in the transition from land to sea (9-18 mya) (Crowe-Riddell et al., 2016; Sanders et al., 2008).

Compared to the flat mechanoreceptors of their terrestrial hydrophiine relatives, scale mechanoreceptors in sea snakes protrude further from the skin surface and form a smooth dome-shaped structure (SD) (Crowe-Riddell et al., 2016; Povel & Van der Kooij, 1997). This SD organ may enhance mechanoreception in sea snakes by increasing the surface area over which water can flow. We hypothesise that enlarged SD mechanoreceptors would allow sea snakes to detect water motion to perceive the movements of potential predators and prey, or turbulence deflected from nearby environmental features (Crowe-Riddell et al., 2016; Crowe-Riddell, Williams, et al., 2019). Morphologically similar, and potentially homologous, SD mechanoreceptors have been described in freshwater *Helicops spp.* snakes (Dipsadidae) (García-Cobos et al., 2021; Velasquez-Cañon et al., 2024) that actively forage for fish and amphibians in lakes and/or streams. Some species of sea snakes have independently acquired much higher coverage of mechanoreceptors over their cephalic scales (4-6%) compared to approximately 2% coverage in other sampled terrestrial snakes (Crowe-Riddell et al., 2016), suggesting there are species-specific selection pressures on mechanoreception based on ecology. However, previous studies in sea snakes have been limited to a single cephalic scale due to the difficulty in quantifying such small and numerous scale mechanoreceptors and ensuring repeatability for comparisons across taxa (Crowe-Riddell et al., 2016, 2019). Scale mechanoreceptors are also highly specular under light microscopy (Crowe-Riddell at al., 2016), making it especially difficult to reliably detect the three-dimensional surfaces of the scales over the head.

In this study, we focus on the olive-headed sea snake, *Hydrophis majo*r, an open water predator with a specialised diet of catfish (Plotosidae). This fully marine species has one of the highest values for mechanoreceptor size, coverage and density among all snakes recorded to date (Crowe-Riddell et al., 2016), but the full distribution of mechanoreceptors over the head has not been comprehensively quantified in any sea snake. The aims of this study are to describe mechanoreceptor morphology using scanning electron microscopy and histology, and quantitatively analyse mechanoreceptor traits (number, density, coverage, height) over the head through the novel application of gel-based 3D profilometry. In addition to creating the first density-based heatmaps of scale mechanoreceptor distribution over the head, we discovered a new type of scale mechanoreceptor that is distinct in its shape and potentially its function (compared to the more abundant SD).

## 2. MATERIALS AND METHODS

### 2.1. Specimens

Ten adult *Hydrophis major* were used in this study (Table 1). Nine adult snakes were collected off Exmouth and Dampier, Western Australia, with approval from The University of Adelaide Animal Ethics Committee (Science) (Approval number S/2021-017) and the WA state (collection permit number FO25000393). Snakes were euthanised with a lethal injection into the body cavity of diluted pentobarbitone (6mg/mL) at a dosage of 150 mg/kg using a 23-30G needle. In addition, a single specimen was sourced from the South Australian Museum. Most specimens were initially frozen, then preserved in 10% formalin and stored in 70% ethanol. A subset of skin samples were taken from specimens fixed fresh by immersion in 4% paraformaldehyde in 0.1 M phosphate buffer or Karnovsky’s solution (2% paraformaldehyde and 2.5% glutaraldehyde in 0.1 M phosphate buffer) for histology and scanning electron microscopy. Measurements of the specimen head (length, width, depth), snout to vent length and tail length were recorded using digital callipers or a tape measure (Table 1).

**Table 1:** *Hydrophis major* specimens and methodologies used in the study. Nine specimens were collected in the field (JCR or KLS identification [ID] numbers) and a single specimen (R ID number) was sourced from the South Australian Museum. SVL = snout to vent length, TL = tail length, GelSight = gel-based 3D profilometry, SEM = scanning electron microscopy, LM = light microscopy histology.

| ID number | Sex | SVL (cm) | TL (cm) | Method | Location |
| --- | --- | --- | --- | --- | --- |
| R67292 | NA | NA | NA | Gelsight | Gulf of Carpentaria, QLD |
| JCR0038 | M | NA | NA | Gelsight, SEM, LM | Exmouth, WA |
| KLS1502 | M | 88.2 | 16 | Gelsight, SEM, LM | Exmouth, WA |
| KLS1068 | M | 72.5 | 7.5 | Gelsight | NA |
| KLS1546 | M | NA | NA | Gelsight, SEM, LM | Dampier, WA |
| KLS1562 | F | 110.5 | 12.3 | Gelsight | NA |
| KLS1334 | F | 79 | 10 | Gelsight | Exmouth, WA |
| KLS1337 | M | 104 | 14 | Gelsight | Exmouth, WA |
| KLS1442 | M | 85 | 6.5* | Gelsight | Exmouth, WA |
| KLS1463 | F | 100 | 12.5 | Gelsight | Exmouth, WA |

\* Tail partly removed by a predator

To investigate the ultrastructure and distribution of scale mechanoreceptors, the head skin of three individuals of *Hydrophis major* was removed and nine cephalic scales were dissected for microscopy. Where possible, four scales from the left side of the head were prepared for light microscopy (LM) and five scales from the right side were prepared for scanning electron microscopy (SEM) (Table 1). Note that for certain specimens, only one of the two sets of scales could be collected in which case priority was given to scales prepared for scanning electron microscopy.

### 2.2. Scanning electron microscopy and histology

The ultrastructure of the mechanoreceptors from five scales of *Hydrophis major* (*n* = 3 individuals) were imaged using scanning electron microscopy (SEM): the lower postocular scale, the supralabial scale located beneath the eye, a parietal scale, the rostrum and a nasal scale (Figure 1). Where possible, all scales were dissected from the right side of the head. The scales were rinsed in three changes of 0.1M Sorensen’s buffer, each for 30 minutes and post-fixed in 1% osmium tetroxide (OsO4) in 0.1M Sorensen’s buffer for two hours. The scales were then rinsed in two changes of milliQ water, each for 30 minutes, and dehydrated in a sequential series of ethanols (30%, 50%, 70%, 80%, 90%, 95% and 100%), each for 30 minutes, and critical point dried for 1.5 hours using a Quorum E3000 Critical Point Dryer (Quorum Technologies Ltd., Laughton, East Sussex, UK). Samples were sputter coated with a Polaron SC7640 Sputter Coater with platinum target (Polaron, Watford, UK) for approximately 35 seconds. The samples were imaged with a Hitachi Ultra-High-Resolution Schottky Scanning Electron Microscope (model SU7000, Hitachi, Tokyo, Japan) in high vacuum mode with accelerating voltages of between 3kV and 7kV, and a spot size of 30. Digital images were taken using the lower, upper and middle detectors. Additionally, navigation images of each of the analysed scales were taken by stitching together low magnification images of the surface of each scale.

**Figure 1:**
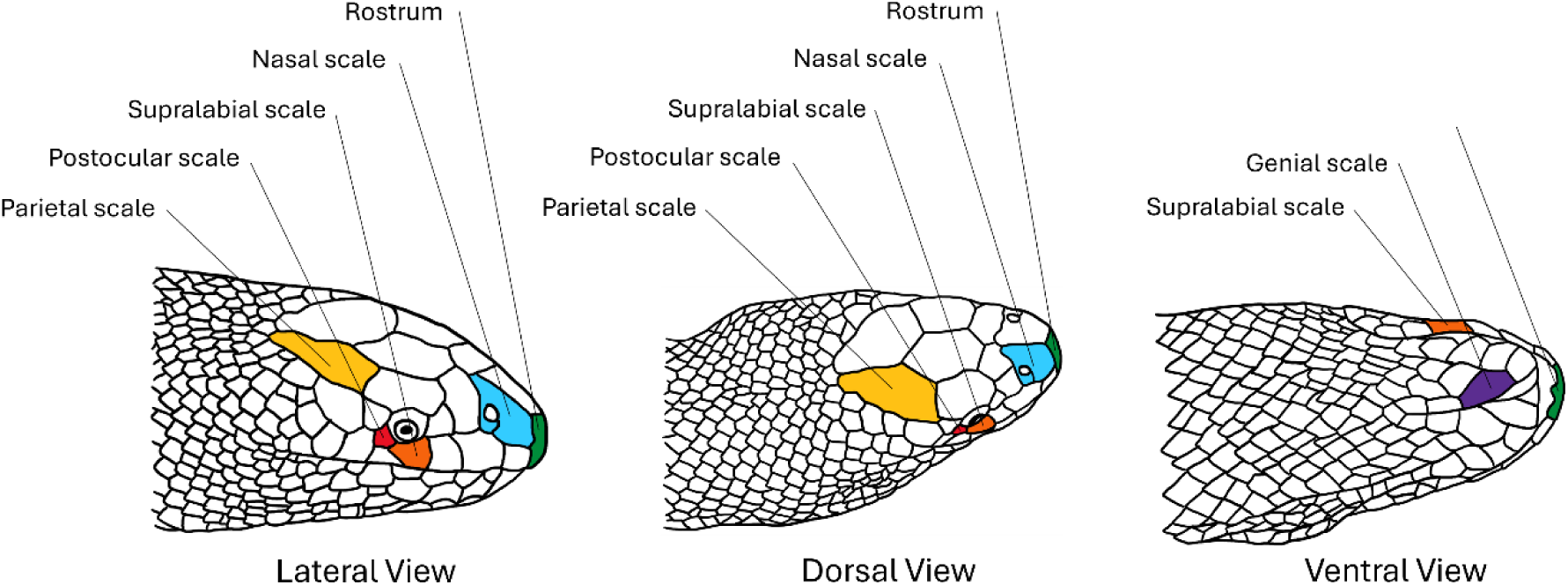
Schematic of the head of an adult *Hydrophis major* Indicating the location of six cephalic scales sampled in this study.

SEM images were measured using the Fiji open-source platform for biological-image analysis (Schindelin et al., 2012), software version 1.8.0_332 (64-bit Java, Image J 1.54). The images enabled the different mechanoreceptor types to be characterised, along with their location on each respective scale and viewing angle. Where possible, measurements of the height, outer diameter, inner diameter and the distance between receptors (both centre to centre and edge to edge) were also recorded. Any damaged mechanoreceptors were not used in any analysis.

The underlying structure of the scale mechanoreceptors of *Hydrophis major* (*n* = 3 individuals) was investigated using histology and light microscopy. Four scales were dissected from the left side of the head where possible: the lower postocular scale, the supralabial scale located beneath the eye, a parietal scale and a nasal scale (Figure 1). The scales were rinsed in three changes of 0.1M Sorensen’s buffer, each for 30 minutes and post-fixed in 1% osmium tetroxide (OsO4) in 0.1M Sorensen’s buffer for two hours. The scales were then rinsed in two changes of milliQ water, each for 30 minutes, and dehydrated in a sequential series of ethanols (30%, 50%, 70%, 80%, 90%, 95% and 100%), sequentially embedded in Spurr’s resin (25%, 50%, 75%, 100%), placed under vacuum for 1 h and then dried in an oven at 60°C for three days. The scales were sectioned to obtain 0.75 μm cross sections using a Leica Ultramicrotome EM UC6 (Leica Microsystems Pty Ltd, Macquarie Park, NSW, Australia) fitted with a glass knife. The sections were mounted onto slides, stained with Toluidine blue and scanned with a Zeiss Axioscan 7 slide scanner (Carl Zeiss AG, Jena, Germany) using brightfield illumination at 20x magnification. The images were analysed and exported using the Zeiss Zen software (version 3.9). The thickness of the Oberhäutchen, stratum corneum and stratum germinativum and the diameter of dermal papilla were measured using a micrometer tool.

### 2.3. Gel-based stereo profilometry

#### 2.3.1. Capturing scale mechanoreceptor topography

A GelSight 1.0X Mobile Core (GelSight Inc., Waltham, Massachusetts, USA) was used to capture a three-dimensional profile of the surface topography of six cephalic scales of *Hydrophis major* (*n* = 10 individuals). Cephalic scales were sampled on the lateral, ventral, dorsal and frontal planes of the head (Figure 1): parietal, nasal, lower postocular scale, supralabial scale located beneath the eye, rostrum and second genial scale. Additionally, for one specimen (R67292), the surface topography of the entire head was captured to create a heatmap of the density and distribution of scale mechanoreceptors. Specimens had either been fixed in 10% formalin and stored in 70% ethanol or had been previously frozen and measured upon thawing. Surface topography models were analysed using the GelSight software (version 3.1.156.0). GelSight is a non-destructive technology that captures high resolution surface information from any surface, including biological samples, over an area of 8.508 × 7.099 mm, using gel-based profilometry.

#### 2.3.2. Image analyses to quantify scale mechanoreceptors

The 3D topological maps generated from the GelSight scanner were analysed with the surface analysis software MountainsMap version 10 (Digital Surf, France). All 3D surface models of scales were treated with a series of operations to standardise comparisons across samples. To flatten the curved scale, while keeping the surface structures intact, the 3D models were treated with a levelling tool using the least squares plane method. This removes the multiplane form and/or separates the roughness and waviness components of the scale (i.e. removes the global slope or form of the surface). The flattened scale is then represented in a 3D model (7% vertical exaggeration), where the mechanoreceptors are clearly visible as peaks. The surface area of each scale was isolated from the model, removing all surrounding background and/or adjacent scales. Profiles were traced through the midline of a sample of mechanoreceptors to obtain information about their 2D shape.

The Gesight scanner measures depressions in a gel created by the surface topography of an object, where the z-axis is negative and ranges from a z-maximum (the lowest point on the sample) to a z-minimum (the tallest point on the sample). To obtain an accurate value for the height of each scale mechanoreceptor, or indeed any feature on the scale, z-maximum needs to be subtracted from all height measurements. The height of mechanoreceptors on each scale was obtained by subtracting the z-minimum from the z-maximum. Height measurements are, therefore, always given in relation to the lowest point on the scale.

To automatically detect mechanoreceptors on a given sample, a particle analysis was conducted using the threshold segmentation tool, which separates any area lying above a certain threshold from the background area, creating isolated particles. Thresholds were manually set for each sample as the smoothing of the scale does not flatten to the same level on all scales. Based on SEM, different classes of mechanoreceptors were identified and distinguished in the surface models using height and roundness thresholding (Supplementary Table 1). The following measurements are obtained for each sample: number of mechanoreceptors, numerical density (density, mm²), percentage of scale occupied by mechanoreceptors (coverage, %) and range and mean height of mechanoreceptors (height, μm).

To assess for potential impacts of specimen storage (either ethanol or frozen) on mechanoreceptor traits, two specimens were measured with the GelSight scanner prior (thawed) and post fixation in 10% formalin (stored in 70% ethanol). The four mechanoreceptor traits (number, height, density and coverage) were compared for the same scale under the two storage conditions with a t-test. Finally, the accuracy of the GelSight scanner surface topography to capture the distribution of scale mechanoreceptors was assessed for two scales (supralabial and postocular) by comparing images captured via GelSight with SEM images on the same scale.

To examine scale mechanoreceptor distribution over the head, a bespoke script was written within Fiji (Schindelin et al., 2012). Firstly, images obtained from 3D profilometry data of a male specimen of *H. major* were manually stitched together and then used to create a sketched outline of the animal. In this image, scale mechanoreceptors were manually identified and marked. The resulting sketched image was loaded into Fiji and the area was broken up into tiles of equal size (1 mm^2^) using regions of interest (ROIs) and marked receptor positions were detected using the "find maxima" function. The number of detected receptors were automatically counted within each tile and subsequently assigned a colour based on the total number counted within.

#### 2.3.3. Statistical analysis

The 3D surface models of scales yielded data on each mechanoreceptor type and on distribution metrics of those receptors over specific scales (height, number, density and coverage). Linear regression on all four mechanoreceptor traits against each individual’s head length were conducted to compare the parameters between scales and the sex of each individual. A non-parametric Kruskal–Wallis test by ranks followed by a Dunn’s test with Bonferroni correction for p-values was conducted to test for the patterns detected by the linear regression and further investigate whether scale and/or sex was associated with mechanoreceptor traits. Several tests were applied to mechanoreceptor traits acquired from the gel-based 3D profilometry including an analysis of variance (ANOVA). In the case where the data is distributed normally and has equal variance, a non-parametric Kruskal–Wallis test by ranks was used. In the case where the data did not follow the assumptions necessary for an analysis of variance, a Dunn’s test followed with Bonferroni correction for p-values. All statistical tests and data visualisation were conducted using the R software v.4.3.3 (R Core Team, 2024) and the following packages: ggplot2 v.3.5.0 (Wickham, 2009), readxl v.1.4.3 (Wickham, Bryan, et al., 2023), dplyr v.1.1.4 (Wickham, François, et al., 2023), FSA v.0.9.5 (Ogle et al., 2023), broom v.1.0.5 (Robinson et al., 2024), tidyverse v.2.0.0 (Wickham et al., 2019), nlme v.3.1.164 (Pinheiro et al., 2024; Pinheiro & Bates, 2000), ggthemes v.5.1.0 (Arnold et al., 2024) and patchwork v.1.2.0 (Pedersen, 2024).

## 3. RESULTS

Fixation and ethanol preservation of the specimen did not significantly impact the scale mechanoreceptor traits of height (t = 0.88, df = 15.74, p-value = 0.39), number (t = 0.49, df = 15.96, p-value = 0.63), density (t = 0.47, df = 15.59, p-value = 0.64) or coverage (t = 1.35, df = 11.99, p-value = 0.20). Thus, whether specimens were fixed or frozen was not considered in subsequent analyses.

### 3.1. Scale mechanoreceptor characterisation

Scanning electron microscopy (SEM) and histology revealed two types of scale mechanoreceptors that differed in their outer surface morphology: symmetrical smooth domes versus asymmetrical pointed peaks, which could be differentiated using gel-based 3D profilometry (Figure 3, Figure 2, Supplementary figure 1).

**Figure 2:**
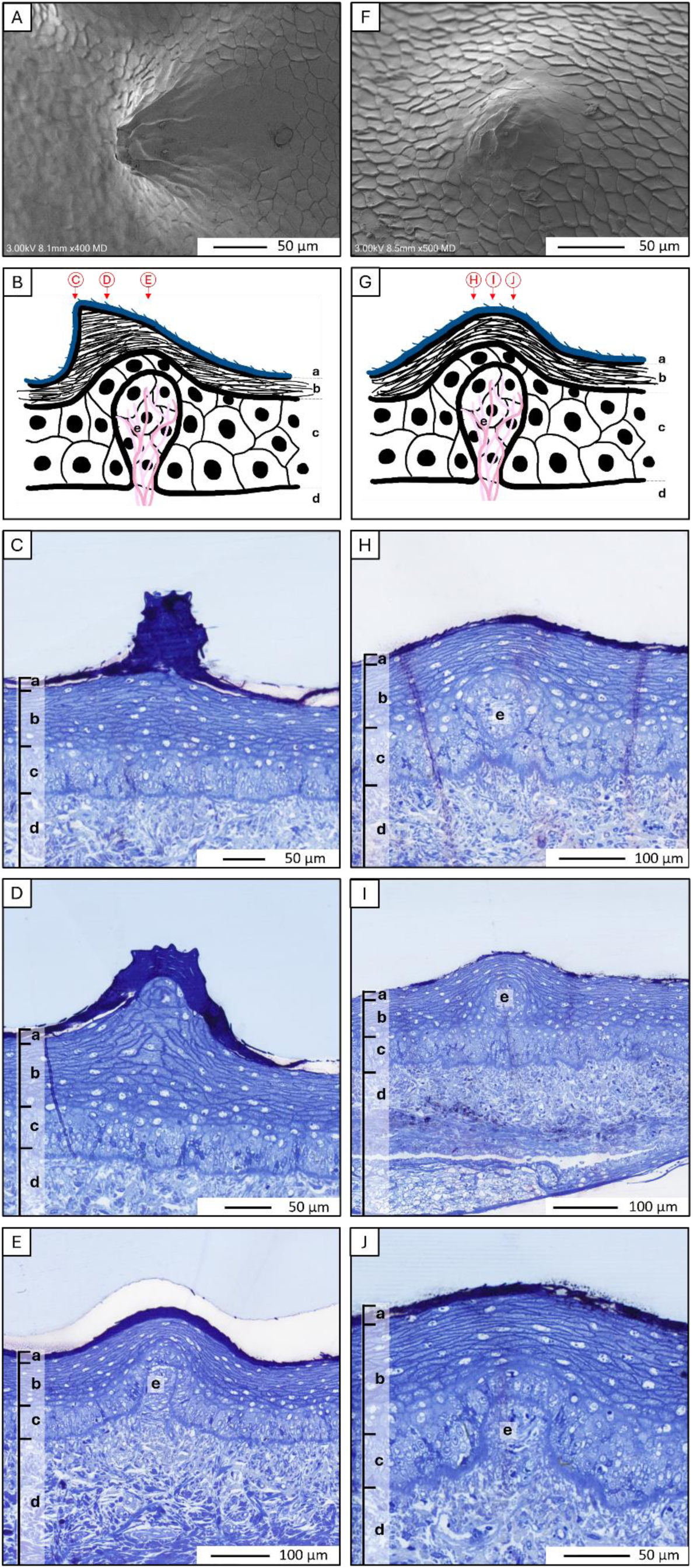
Two types of cutaneous mechanoreceptors in *Hydrophis major*: the asymmetrical peak type is represented in the first column and the smooth dome type is represented in the second column. A) and F) are scanning electron microscope images captured by the middle detector. B) and G) are schematic representations for each mechanoreceptor type and show outer shape and skin layers. There are the three layers of the epidermis: Oberhäutchen (a), stratum corneum (b) and stratum germinativum (c); as well as the dermis (d) and dermal papilla (e). C), D), E), H), I) and J), are light micrographs of transverse sections stained with Toluidine blue corresponding with red markers in the schematic diagram.

**Figure 3:**
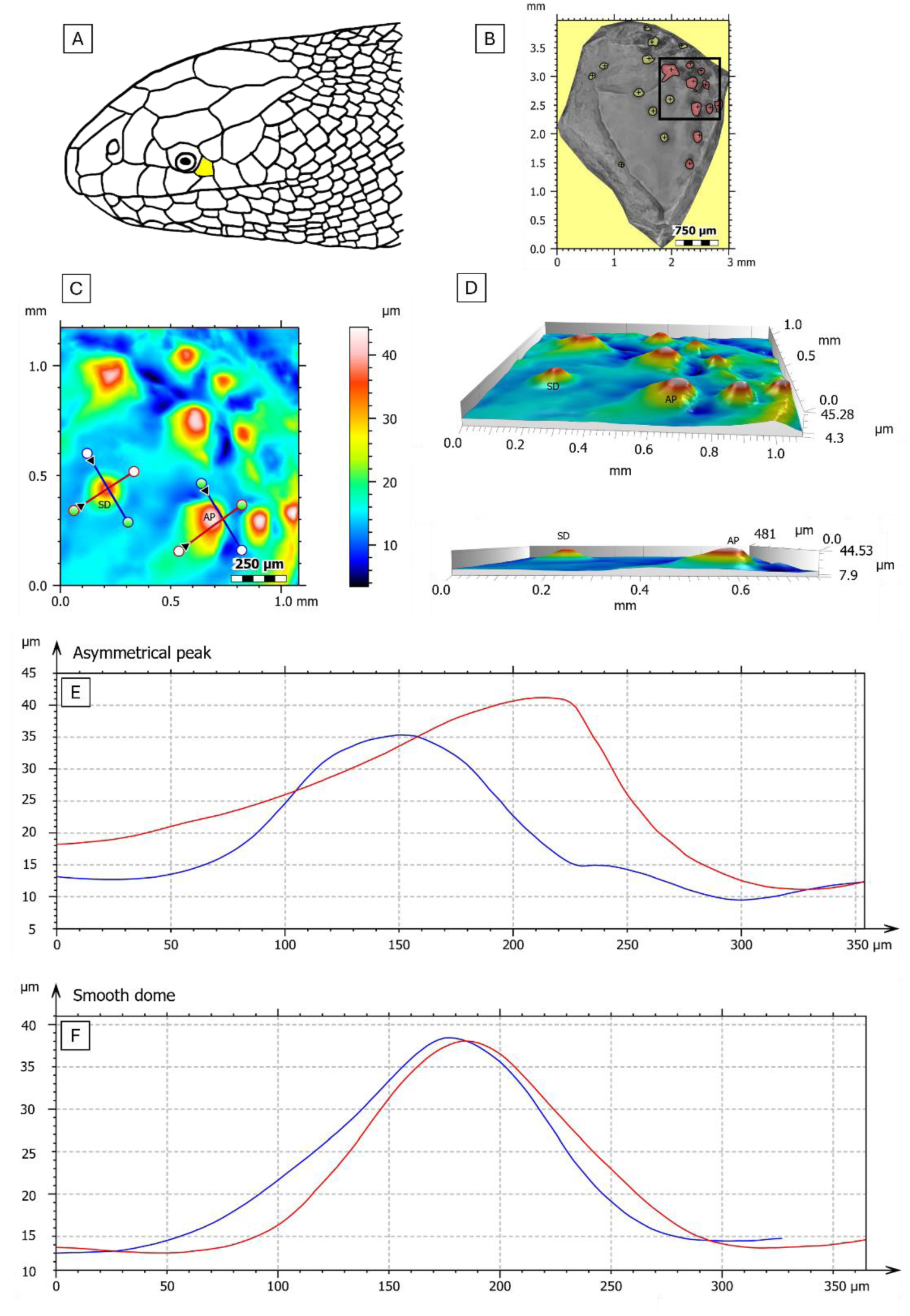
Comparison of the height and shape of two mechanoreceptor types on the postocular scale of *H. major* using gel-based 3D profilometry in Mountains Map software. A) Location of the postocular scale. B) Particle detection of smooth dome (SD in yellow) and asymmetrical peak (AP in red) mechanoreceptors. C) Extracted area as indicated by the black square in (B) showing SD and AP mechanoreceptors. Colour gradient indicates the height (μm) of the scale. Crosshair lines correspond cross sectional profiles in (F, G). (D) 3D model of the extracted area. (E) Lateral aspect of the extracted area that shows the height and degree of roundness of SD and AP mechanoreceptors. (F-G) cross-sectional height profiles of SD and AP mechanoreceptors. Red lines correspond to rostral-caudal sections and blue lines correspond to dorso-ventral sections.

The most commonly encountered mechanoreceptor type is a smooth dome. This type of scale mechanoreceptor was found on every scale investigated. It is a dome-shaped structure (Figure 2) characterised by an unbroken surface layer of cells, the Oberhäutchen, which forms a continuous elevation in the skin. The term smooth refers to the continuous nature of the surface of the scale mechanoreceptor, not from the fact that the surface is perfectly rounded. Various stages of ageing and deformation of these smooth domes were identified (Supplementary figure 1). Some are scraped domes, where the Oberhäutchen appears to have been scraped off or peeled back at the centre of the dome. Some are slightly hollowed domes, where there is a caldera-like hole in the middle of the dome, revealing the central cell cluster, originating from the dermal papilla. Some domes were flattened where the dome appeared to have experienced a mechanical force that has compacted it and flattened out the top of the dome.

The second, rarer, type of scale mechanoreceptor has an asymmetrical peak. These mechanoreceptors are large and pointed, resembling the traditional shape of a mountain peak, except that the tip may be slightly off-centre (Figure 3), giving the mechanoreceptor an almost thorn-like appearance (in between the shape of a right triangle and a scalene triangle). Under SEM, the outer layers appear as spikes formed by the individual epithelial cells of the Oberhäutchen at the top of the peak (Figure 2).

The two types of mechanoreceptors differ in shape and height and could be differentiated using SEM (Supplementary figure 1) and/or the 3D profiles generated by the GelSight scanner (Figure 3). According to the measurements taken from SEM images, the asymmetrical peaks have a diameter of 139.72 μm (± 52.31 μm), which never falls below 100 μm, while smooth domes have a mean diameter of 137.48 μm (± 73.09 μm). SEM measurements of the two mechanoreceptor types were not significantly different in diameter (chi-squared = 0.76, df = 2, p-value = 0.68) or height (chi-squared = 3.11, df = 2, p-value = 0.05). The angle of the detector to the scale in the scanning electron microscope, however, limited the number of measurements of the mechanoreceptor height that could be taken accurately. In contrast, the GelSight scanner revealed that smooth domes and asymmetrical peaks are significantly different in height (chi-squared = 77.006, df = 1, p-value = < 2.2e^-16^). Smooth domes have a mean height of 31.88 ± 19.03 while asymmetrical peaks have a mean height of 43.65 ± 22.57.

Measurements taken from SEM measurements showed that asymmetrical peaks, smooth domes and damaged domes did not differ significantly in their height when analysing all scales together, although the lack of significance is likely due to the extremely low number of asymmetrical peaks that could be measured accurately (n = 3), as clear height differences were detected with the GelSight scanner.

Histology under light microscopy on postocular and supralabial scales revealed dermal papilla underlying both scale mechanoreceptor types. Multiple sections were taken in regions without a mechanoreceptor, through a smooth dome and through an asymmetrical peak (Figure 2). On both samples, four layers of the skin are clearly distinguishable. A scale mechanoreceptor appears as a rounded elevation in the surface of the skin and is recognisable as a sensory organ by the presence of a dermal papilla. It is a local protrusion of the dermis (skin layer containing living cells, collagen fibres, nerve bundles, blood vessels and glandular tissue) into the epidermis (three outer layers of the skin) forming a cluster of cells and resulting in the compression and consequent thinning of the upper epidermal layers (Table 2).

**Table 2:** Thickness measurements of three epidermal layers of *Hydrophis major* for both postocular and supralabial scales. Each measurement was taken in four different locations on the scale: in an area without a mechanoreceptor (flat skin), above the dermal papilla of a smooth dome, above the dermal papilla of an asymmetrical peak and at the highest point of an asymmetrical peak.

| Skin layer | Asymmetrical peak<br>highest point (µm) | Asymmetrical peak<br>dermal papilla (µm) | Smooth dome<br>dermal papilla (µm) | Flat skin (µm) |
| --- | --- | --- | --- | --- |
| Oberhäutchen | 34.56 ± 2.59 | 13.36 ± 0.16 | 12.85 ± 4.23 | 6.09 ± 0.81 |
| Stratum corneum | 125.67 ± 1.06 | 34.98 ± 0.54 | 41.32 ± 5.89 | 60.05 ± 6.30 |
| Stratum germinativum | 48.09 ± 4.93 | 16.59 ± 5.97 | 16.93 ± 5.25 | 39.87 ± 6.12 |

Asymmetrical peak mechanoreceptors are primarily composed of a thickened outer layer of the epidermis including the Oberhäutchen (outermost layer of highly cornified cells of the epidermis, dark blue in Figure 2), stratum corneum (dead, flattened, strongly keratinized and tightly joined cells that lack a nucleus) and stratum germinativum (a single layer of large vertically elongated cells). In contrast, the smooth domes are composed of thinner layers of the epidermis including Oberhäutchen, stratum corneum and stratum germinativum. In the case of an asymmetrical peak, the dermal papilla is not located subjacent to the peak but rather several micrometres displaced under the gently sloping side of the thorn-shaped mechanoreceptor (Figure 2). The epidermis above the dermal papilla of the asymmetrical peak has a similar dermal thickness to the dermal papilla of the smooth domes.

The Oberhäutchen was uniform in thickness across the scale except at the highest point of the asymmetrical peak mechanoreceptors (Figure 2, C and D, Table 2) where it is 2.6 times thicker (34.56 μm ± 2.59 μm c.f. 13.06 ± 8.92 μm of flat skin; chi-squared = 13.47, df = 3, p-value = 0.004). No significant differences were found between the investigated scale types (chi-squared = 4.5113, df = 1, p-value = 0.034) or across individuals (chi-squared = 7.5236, df = 2, p-value = 0.03367).

Similar to the analysis of the Oberhäutchen, the thickness of the stratum corneum shows no significant difference between scale types (chi-squared = 3.61, df = 1, p-value = 0.060) or across individuals (chi-squared = 5.14, df = 2, p-value = 0.057). A Kruskal-Wallis test (chi-squared = 14.65, df = 3, p-value = 0.002) followed by a Dunn’s test using the Bonferroni correction for p-values identified patterns in the thickness of the layer depending on the location on the scale. The stratum corneum is similar across different regions including a smooth dome mechanoreceptor, an area of flat skin with no mechanoreceptor and the area above the dermal papilla in an asymmetrical peak mechanoreceptor (Figure 2). In these regions, the thickness of the stratum corneum has a mean of approximately 55.45 μm (± 26.97 μm), but is significantly thinner where it overlies a dermal papilla (c.f. smooth dome mechanoreceptors to flat skin, Z = 2.64, p-adj = 0.050, and superior to the dermal papilla of an asymmetrical peak mechanoreceptor c.f. flat skin , Z = -2.50, p-adj. = 0.074). However, at the apex of an asymmetrical peak mechanoreceptor, i.e. not directly superior to the dermal papilla, the stratum corneum has a thickness of 125.67 μm (± 1.1 μm), which is 2.3 times thicker than in any other area of the scale (Figure 2 D, Table 2). When counting the number of cell layers comprising the stratum corneum, there is a clear difference between any mechanoreceptor associated area, where there are between 10 and 13 cell layers, and an area lacking mechanoreceptors where there are 16 layers.

In contrast to the Oberhäutchen and the stratum corneum, a Kruskal-Wallis test reveals significant differences in the thickness of the stratum germinativum between scales (chi-squared = 8.41, df = 1, p-value = 0.004) and individuals (chi-squared = 9.66, df = 2, p-value = 0.008). An ANOVA (F-statistic = 32.14, df = 3, p-value = 8.958e^-07^) followed by a Dunn’s test using the Bonferroni correction for p-values shows that the thickness of the stratum germinativum superior to the dermal papilla is is similar between a smooth dome mechanoreceptor and an asymmetrical peak mechanoreceptor (Table 2), with a mean of 16.87 μm (± 5.1 μm). However, the stratum germinativum is thicker in areas without a mechanoreceptor, where it reaches a thickness of 39.87 μm (± 6.1 μm), and at the highest point of an asymmetrical peak, where it reaches 48.09 μm (± 4.9 μm). Therefore, above any given dermal papilla, irrespective of the mechanoreceptor type, the stratum germinativum is 2.6 times thinner.

For the dermal papilla, a Kruskal-Wallis test and an analysis of variance reveals that there are no significant differences in its diameter between scales (chi-squared = 1.6364, df = 1, p-value = 0.201), across individuals (chi-squared = 1.6364, df = 1, p-value = 0.201)or between location on the scale (F-statistic = 0.4054, df = 1, p-value = 0.542). For both mechanoreceptor types, the dermal papilla is a rounded protrusion of the dermis and has a mean diameter of 47.74 ± 8.1 μm (Table 2) and is composed of five to eight cells.

### 3.2. Distribution of scale mechanoreceptor types

Images taken of the same cephalic scales by the GelSight and under SEM show no change in the number or distribution of mechanoreceptors (Supplementary Figure 2). Therefore, SEM and gel-based 3D profilometry using a GelSight scanner are complementary techniques that can be used interchangeably to map the distribution of mechanoreceptors.

Distribution of the two mechanoreceptor types was observed under SEM and compared to gel-based 3D profilometry from the GelSight scanner. While smooth dome mechanoreceptors were found on all scales, asymmetrical peak mechanoreceptors were found only on postocular, parietal, supralabial and (in one individual) a nasal scale. Irrespective of mechanoreceptor type, SEM also revealed that scale mechanoreceptors are generally clustered more closely (have a higher density) on the postocular and supralabial scales compared to any other scales, although it was not possible to examine rostrum or genial scale due to damage of those scales in the observed specimens. SEM analyses of the differentiation between mechanoreceptor types and their distribution were corroborated using the GelSight scanner and confirmed that asymmetrical peaks were detected only on the supralabial, nasal, postocular and parietal scales (Figure 4).

**Figure 4:**
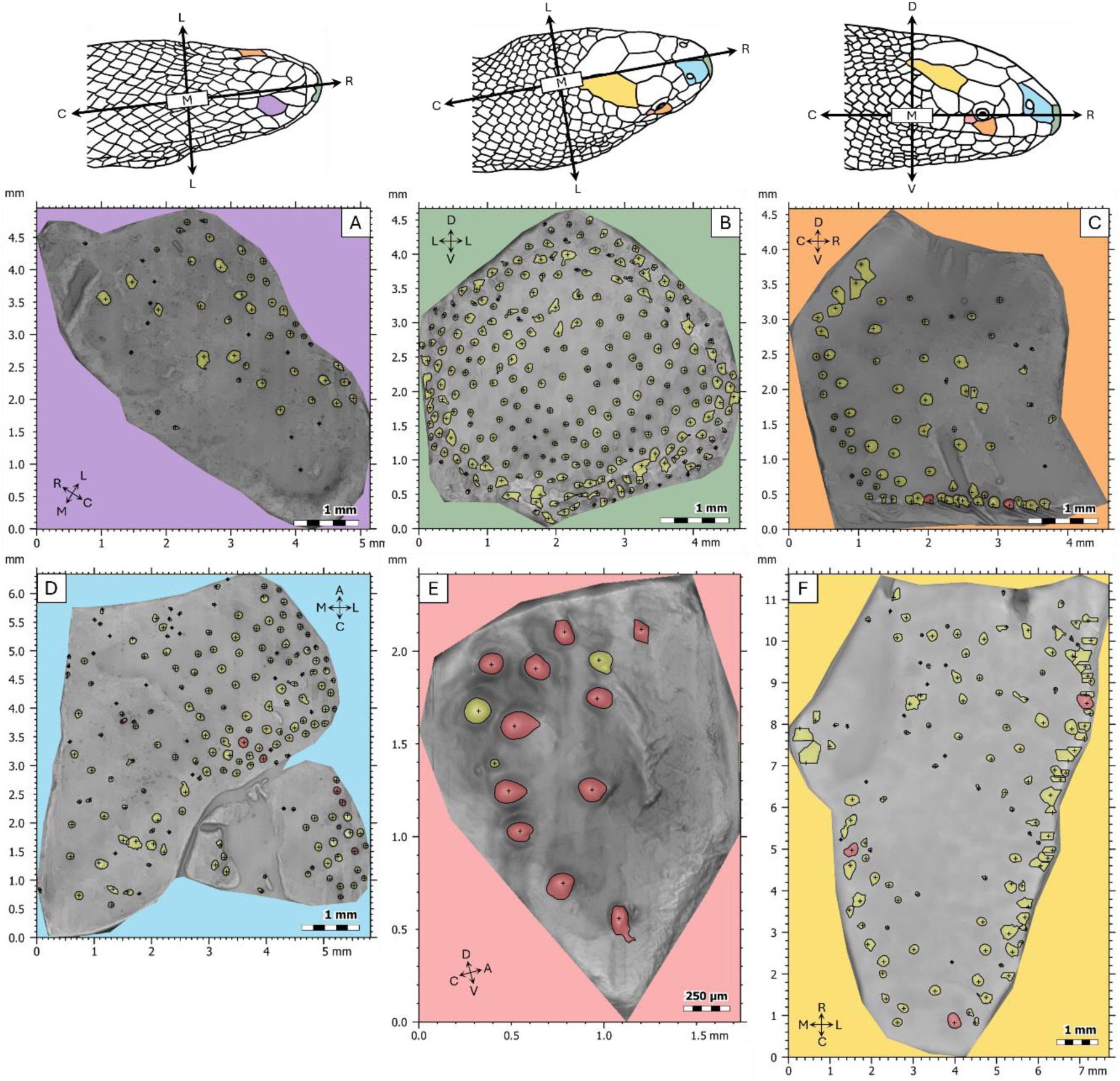
Distribution of smooth dome mechanoreceptors (yellow dots) and asymmetrical peak mechanoreceptors (red dots) on cephalic scales in *Hydrophis major* using gel-based 3D profilometry. The orientation labels are C for caudal, R for rostral, L for lateral, M for medial, D for dorsal and V for ventral. The colours on the snake diagrams correspond to the background colour of each panel. The scales shown are the genial scale in ventral view (A), the rostrum in frontal view (B), the supralabial scale in lateral view(C), the nasal scale in dorsal view (D), the postocular scale in lateral view (E) and the parietal scale in dorsal view (F). All scales were taken from the right side of the body except the genial scale.

Irrespective of mechanoreceptor type, Figure 4 shows a higher abundance of scale mechanoreceptors close to the scale edges. It is also evident that dorsal and ventral scales, such as the observed nasal, parietal and genial scales, show a preferential alignment of scale mechanoreceptors toward the lateral edge of the scales (Figure 4, A, D and F). Lateral scales, such as the postocular and the supralabial scales, show a higher density toward the caudal edge of the scale (Figure 4, C and E). Note that, in addition to a caudal alignment, supralabial scales also show an alignment on the ventral scale edge. Mechanoreceptors on the rostral scale (Figure 4 B) do not show any preference for a particular scale edge.

Asymmetrical peak mechanoreceptors are relatively rare (1-19 per scale), while smooth dome mechanoreceptors are at least 20 times more frequent on the supralabial scales (3-426 per scale), and 30 times more frequent on the nasal scales (Figure 5). Asymmetrical peak mechanoreceptors were entirely absent on certain scales such as the genial scale and the rostrum. However, the one exception to this pattern is the postocular scale, where asymmetrical peak mechanoreceptors are more numerous than smooth dome mechanoreceptors (7.5 ± 5.46 smooth domes and 15.1 ± 8.16 asymmetrical peaks). Male and female *H. major* typically have similar numbers of asymmetrical peak mechanoreceptors on their scales, but females possess higher numbers of smooth dome mechanoreceptors than males (Supplementary figure 3). Note that this level of sexual dimorphism must be taken with an element of caution as the sample number was relatively low and not all specimens had all scales intact and that were usable for comparison (n = 10 with 6 males, 3 females and one unknown).

**Figure 5:**
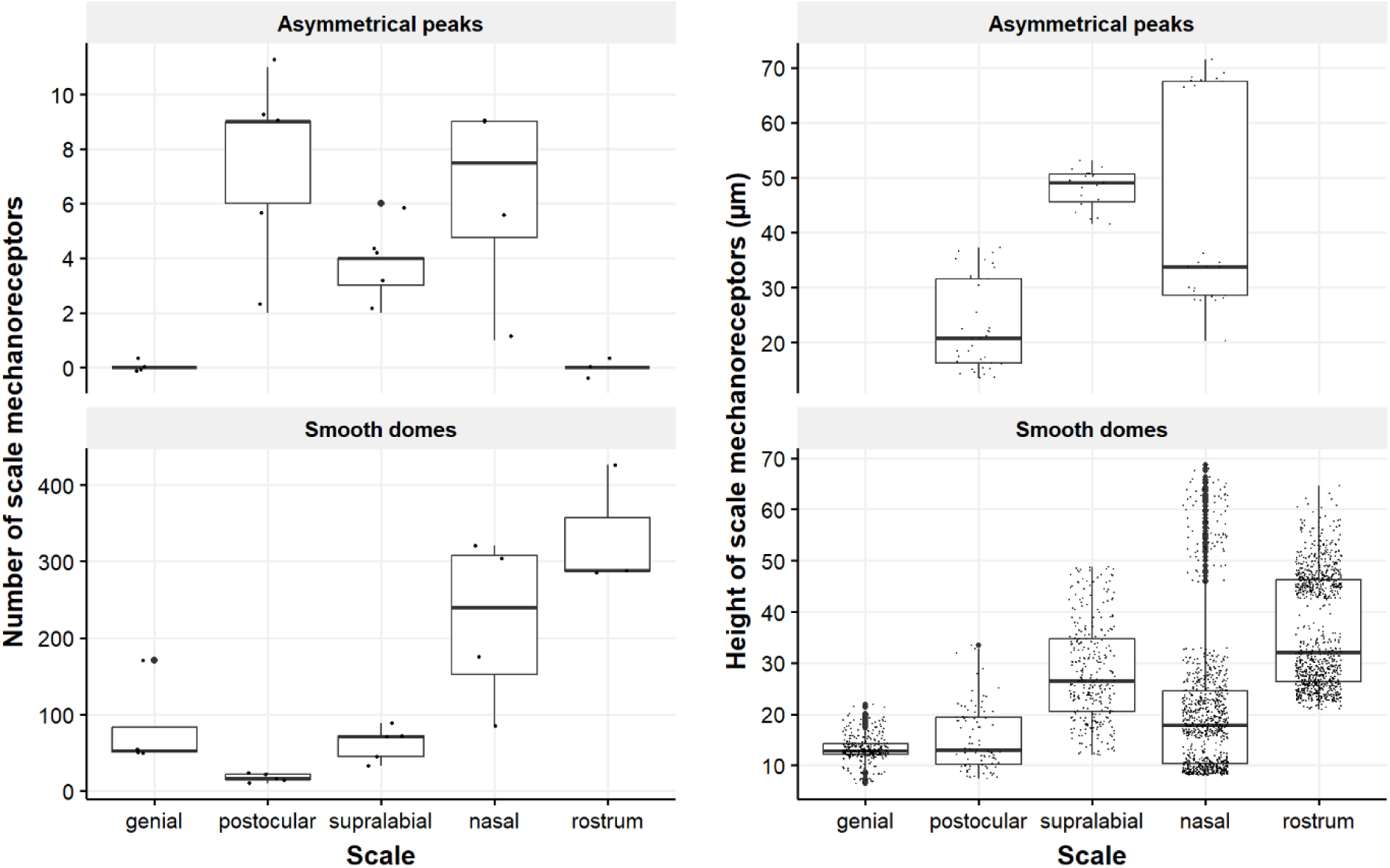
Boxplot of the number and height (µm) of the two types of mechanoreceptors (smooth dome and asymmetric peak) per scale in *Hydrophis major* across five cephalic scales. Each box represents the interquartile range (IQR), with the line inside the box marking the median. Outliers are presented as large points. Statistical analysis using the Kruskal-Wallis test revealed significant differences in the number of mechanoreceptors across scales (chi-squared = 3118.4, df = 4, p-value < 2.2e^-16^). For the number of mechanoreceptors, the points represent the total number of receptors per individual per scale; note the different y axes. For the height of mechanoreceptors, the points represent individual height for each receptor detected. Statistical analysis using the Kruskal-Wallis test revealed significant differences in the mechanoreceptor height across scales (chi-squared = 1218.2, df = 4, p-value 2.2e^-16^).

When comparing smooth dome and asymmetrical peak mechanoreceptor types, an overlap in height of mechanoreceptors was revealed across all scales. However, the overlap was essentially eliminated when comparing both mechanoreceptor types for each individual scale (Figure 5), where there is a negative relationship between mechanoreceptor height and head length between scales (Table 3). All mechanoreceptor traits were significantly different between sexes across scales. The level of this significance was further investigated with a Kruskall-Wallis test followed by a Dunn’s test with Bonferroni correction for p-values, which revealed that the observed significance is found between all scale types, with no two scales showing similar patterns in height. For example, looking at the nasal scale, asymmetrical peak mechanoreceptors possess a mean height of 34 µm, while smooth dome mechanoreceptors possess a mean height of 19 µm. Note that, when comparing mechanoreceptor height between sexes, the differences in height become even more pronounced. The same pattern can be found on all the other scales examined. While females seem to have an overall higher number of mechanoreceptors, males have taller mechanoreceptors of both types, with the exception of the genial scale where the mechanoreceptor height is similar to females, and the rostrum where the pattern is inverted (Supplementary figure 3). Note that an ANOVA revealed that there is no significant difference in head length between males and females (F_(1,5)_ = 0.07, p = 0.80) and thus observed differences between sexes are not driven by head size.

**Table 3:** Allometric relationship between head length (HL) and mean height of scale mechanoreceptors, number of scale mechanoreceptors, overall coverage of scale mechanoreceptor and numerical density of scale mechanoreceptor across six scales between males and females. Equations are in the form y = ax^b^, where y is the trait of scale mechanoreceptor, a is the coefficient (elevation), b is the exponent (slope) and x is the head length.

| Traits of sensilla | $x$ | Coefficient, $a$ | Exponent, $b$ | 95% CI | $r^2$ | d.f. | $F$ |
| --- | --- | --- | --- | --- | --- | --- | --- |
| Height ($\mu\text{m}$) | HL | 3.71 | -0.09 | $\pm 0.04$ | 0.01 | 2 | 12.59 |
| Number | HL | 8.93 | -0.19 | $\pm 0.05$ | 0.02 | 2 | 38.3 |
| Coverage (%) | HL | 5.06 | 0.09 | $\pm 0.04$ | 0.06 | 2 | 129.1 |
| Density ($\text{mm}^2$) | HL | 12.74 | -0.16 | $\pm 0.03$ | 0.22 | 2 | 527.4 |

### 3.3. Distribution of all scale mechanoreceptors

The density of all mechanoreceptors (both smooth dome and asymmetrical peak types combined) revealed no significant differences between sexes across all scales (chi-squared = 0.08, df = 1, p-value = 0.78). There were significant differences when comparing coverage and density between scales (Supplementary figure 3). With respect to mechanoreceptor density, comparisons between the genial and supralabial scales and the supralabial and postocular scales were the only comparisons that did not show a significant difference, with all other interscale comparisons showing significant differences (Supplementary figure 4). The scale with the highest density overall was the rostrum, with a mean of 11 receptors per mm². With respect to mechanoreceptor coverage, interscale comparisons showed a significant difference in mechanoreceptor coverage among all scales except the parietal and supralabial scales. The scale with the lowest coverage of mechanoreceptors was the genial scale with less than 2% coverage, while the scale with the highest coverage was the rostrum with a coverage of around 9% (Supplementary figure 4). Density and coverage follow roughly the same pattern.

There is a marked heterogeneity in both coverage and density of mechanoreceptors over the surface of the scales. The level of heterogeneity is seen most effectively in the supralabial scale, where the mechanoreceptors form an almost horizontal line along the edge of the scale adjacent to the mouth opening (Figure 6). A similar pattern is found in adjacent supralabial scales as well as sublabial scales (Figure 6, B and C). These horizontally aligned scale mechanoreceptors are characterised as smooth dome mechanoreceptors, which occur in high density and are so close to each other that the dermal papilla of each mechanoreceptor is separated by only 100 µm from its neighbour, which corresponds to the diameter of the mechanoreceptor. The lack of separation between mechanoreceptors, indicating that they are morphologically at maximal density, is also noticeable when looking at the Oberhäutchen. Rather than the expected thinning between the mechanoreceptors, as is characteristic of a non-mechanoreceptor area, the space between the mechanoreceptors is so narrow that there is a localized thickening of the Oberhäutchen, which reaches a thickness of around 25 μm, approaching that of the highest peak of a scale mechanoreceptor.

**Figure 6:**
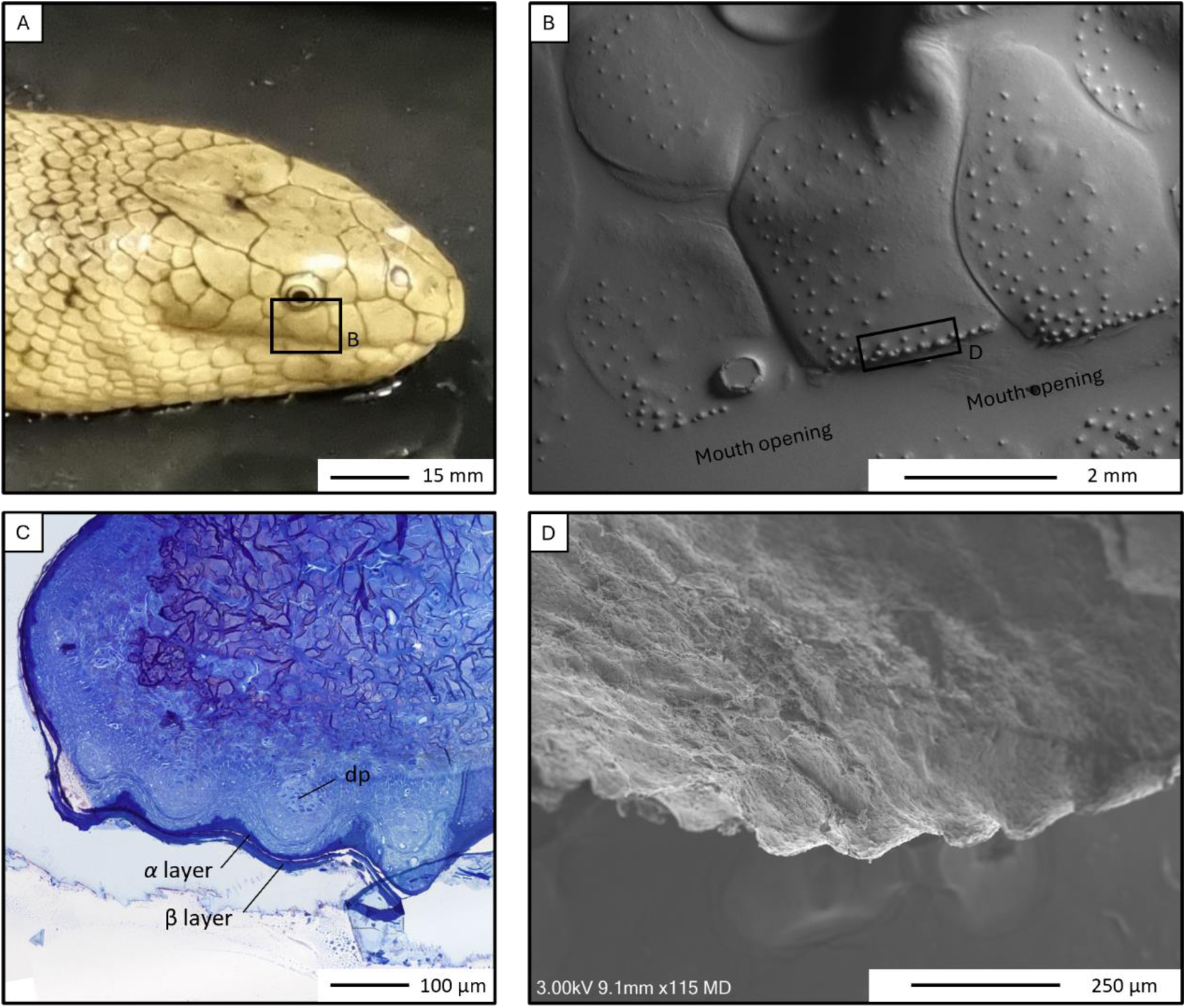
Distribution of smooth dome mechanoreceptors on the fourth supralabial scale of *Hydrophis major*. A) A male *Hydrophis major*. B) GelSight scanner image showing the distribution of cutaneous mechanoreceptors including the alignment of mechanoreceptors along the ventral edge of the scale adjacent to the mouth. C) Light micrograph of a section through a supralabial scale stained with Toluidine blue showing the histology of the mechanoreceptors along its ventral edge, including a dermal papilla (dp). D) Higher power scanning electron micrograph of the horizontal alignment of mechanoreceptors. Image of live snake by Vhon Garcia.

When assessing the total mechanoreceptor distribution over the whole head of *H. major*, the mouth opening exhibits a conspicuous concentration of mechanoreceptors. In Figure 7 (B and D), the highest concentration of mechanoreceptors is found along the lingual edges of the supralabial and sublabial scales, with the highest densities observed on the supralabial scales. Other high-density areas include the rostrum, the anterior end of the nasal scales and supraocular scales (located above the eye). The lowest densities are observed on the ventral side of the head (Figure 7 A) and posterior-dorsal scales towards the body. The distribution of mechanoreceptors within each scale is heterogeneous with local density maxima occurring toward the lateral and caudal edge of the scales, illustrated well in the supralabial scale (Figure 6). Striking examples are the supralabial and postocular (Figure 7 D) and parietal (Figure 7, C) scales. The only exception to this pattern is the rostral scale, which had a more homogenous distribution of mechanoreceptors. With the help of the generated heat map and the individually counted scales, an estimate of 7000 scale receptors were identified over an entire snake head (approximately 700 ventrally, 750 frontally, 1800 laterally and, 1920 dorsally).

**Figure 7:**
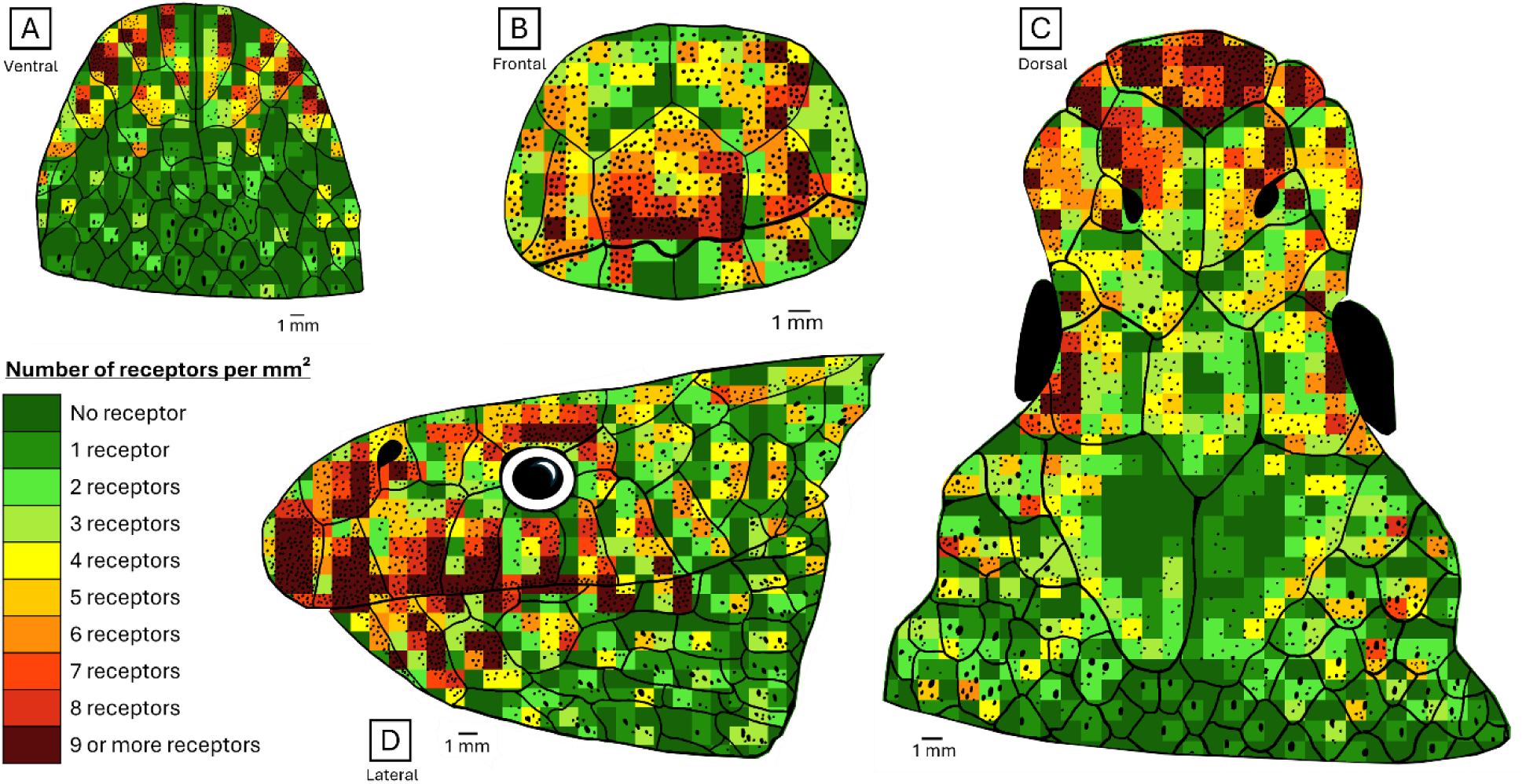
Heatmap of the density and distribution of mechanoreceptors over the head of *Hydrophis major*. The colour gradient represents the density per mm^2^ of scale mechanoreceptors in four views: ventral (A), frontal (B), dorsal (C) and lateral (D).

## 4. DISCUSSION

Previous studies on cephalic mechanoreceptors in sea snakes have focused mainly on the potential for a derived hydrodynamic reception function by comparing the shape and ultrastructure of mechanoreceptors with those in terrestrial species (Crowe-Riddell et al., 2016; Crowe-Riddell, Williams, et al., 2019). This study is the first high resolution quantification of scale mechanoreceptors over the head in any snake and, while previous studies have described the smooth dome type of mechanoreceptors (Crowe-Riddell et al., 2016, 2021), we describe a novel type of mechanoreceptor that we refer to as an “asymmetrical peak”. These two types of scale mechanoreceptors in *Hydrophis major* differ in their morphology and distribution over the head, suggesting different mechanosensory functions. We discuss the potential relevance of the distribution and morphology of these mechanoreceptor types for the foraging behaviour and ecology of *H. major*, as well as the broader significance for hydrodynamic reception in fully aquatic sea snakes.

The novel application of gel-based 3D profilometry to obtain detailed high-resolution models of microscale cephalic skin structures has also yielded comparable results using more traditional and time-consuming methods such as scanning electron microscopy (Supplementary figure 2). The conformation of the gel over the sample creates a high-fidelity topological map, eliminating specular reflections from the scales that can make imaging scale mechanoreceptors in reptiles challenging. In addition to generating data on the number, density and coverage of scale mechanoreceptors, the GelSight scanner replicates the height and shape of scale mechanoreceptors, allowing us to rapidly quantify and differentiate mechanoreceptor types for the first time (Figure 3, 5). Our study supports the use of gel-based profilometry to detect skin structures in a diversity of species (Baeckens et al., 2019; Wainwright et al., 2017), regardless of preservation methods (fixed or frozen) and without significantly affecting the integrity of the specimen.

### 4.1. Morphological characterisation of two scale mechanoreceptor types

We characterised the morphology of the smooth dome (SD) and asymmetrical peak (AP) mechanoreceptor types in *H. major* using both scanning electron microscopy (SEM) and histology. We confirm that both types contain a Meissner-like corpuscle (dermal papilla), homologous to the structure of scale organs described in other species of hydrophiine sea snakes, land snakes (including Hydrophiinae, Colubridae, Leptotyphlopidae) and dipsadid aquatic snakes (Crowe-Riddell, Williams, et al., 2019; García-Cobos et al., 2021; Povel & Van der Kooij, 1997). The dermal papilla and corresponding skin elevation are widely conserved in the scale mechanoreceptors of both land and sea snakes (Crowe-Riddell et al., 2021; García-Cobos et al., 2021; Jackson & Sharawy, 1980; Landmann, 1975). The dermal papilla of AP mechanoreceptors is rostrally located and dissociated from the highest point of the peak (Figure 2). The latter is created by a thickening of all tissue layers but especially of the highly cornified Oberhäutchen. However, the thickness of the epidermal layers above the papilla, as well as the diameter and number of central cells within the dermal papilla, does not vary between the two types of mechanoreceptors.

The two types of mechanoreceptors differ in their surface ultrastructure, size (height and diameter) and their shape (symmetrical and smooth versus asymmetrical and pointed). Most noticeable is the difference in shape, with the SD type being rounded and showing no particular asymmetry or directionality, while the highest point of the thorn-shaped AP points caudally relative to the positioning of the dermal papilla. In addition to their shape, the APs are distinguished by their greater height. Measurements taken from SEM measurements showed that APs, SDs and damaged domes did not differ significantly in their height when analysing all scales together, although the lack of significance is likely due to the extremely low number of APs that could be measured accurately (n = 3), as clear height differences were detected with the GelSight scanner. While there is an overlap between the tallest SD and the smallest AP, the latter are generally taller than the former. Indeed, APs range in height from 13 to 150 μm (mean of 43.65 μm), while SDs have a height between 5 and 137 μm (mean of 31.88 μm), depending on the scale type (Figure 5).

In addition to height, the diameter of the two types of mechanoreceptors also differs. While the mean dimensions of SD scale mechanoreceptors in *H. major* are only slightly larger than what has been observed in a previous study (Crowe-Riddell et al., 2016), our study reveals large variations in the size of the SD type, which range from 30 to 200 μm in diameter, which has not been described previously. These size differences might simply be due to the mechanoreceptors being of different age or the constraints of optimising packing density and thereby sensitivity. Studies on freshwater snakes (*Helicops angulatus, Helicops danieli,* and *Helicops pastazae*) have shown that mechanoreceptor size differs with their location on the animal, and that areas that are more used for mechanoreception have typically larger receptors (Velasquez-Cañon et al., 2024). Others have speculated that the size of the papilla would correspond to the size of the scale elevation (García-Cobos et al., 2021). Although we were not able to test this, our images under SEM clearly show different mechanoreceptor sizes.

Differences in the thickness of each skin layer of the mechanoreceptors may be dependent on the skin shedding cycle. As only epidermal layers are replaced and regenerated during sloughing (Maderson, 1965), one would expect to be able to clearly see the mechanoreceptors throughout the shedding cycle (Jackson & Sharawy, 1980; Landmann, 1975). It is possible that the damaged scale mechanoreceptors observed under SEM may have been different shedding stages of scale mechanoreceptors (Supplementary figure 1). However, how the thickness of the various skin layers with the mechanoreceptors changes in sea snakes is unknown, as their shedding cycle has yet to be fully examined. As many more epidermal layers are present above each mechanoreceptor just before shedding, one would expect their sensitivity to drastically diminish during this period (Povel & Van der Kooij, 1997), which likely influences their temperament and makes them more defensive. This periodic reduction in sensitivity is expected to be even more pronounced in sea snakes than in terrestrial snakes as their skin is often much thicker (Shine et al., 2019) and shed more frequently.

### 4.2. Variation in mechanoreceptor distribution over the head

Variation in mechanoreceptor abundance was found for both mechanoreceptor types and between the different planes of the snake head (Figure 7). The number and density of scale mechanoreceptors consistently decreases from the snout to the caudal end of the head. Indeed, the highest number and overall density of mechanoreceptors was found on the frontal plane of the head with highest number/density on the tip of the snout (Supplementary figures 3 and 4). During locomotion, the front of the snake head encounters and interacts with its environment first and this potentially allows the animal to actively sense hydrodynamic flow for navigation while foraging, avoiding predators and searching for conspecifics. Therefore, the ability to navigate around and into crevices or within the water column would heavily rely on being able to accurately detect mechanosensory information in front of the head with increased acuity. It is thus not surprising to find the highest densities of mechanoreceptors in this region as has been described in both aquatic and terrestrial snakes (García-Cobos et al., 2022; Jackson & Sharawy, 1980). The lowest density of receptors is found on the ventral scales, where the head scales transition to body scales much faster than any of the other planes (Kordi & Shabanipour, 2012). While the SD mechanoreceptors are found on every head scale, the AP mechanoreceptors are restricted only to certain scales: the nasal, postocular, parietal and supralabial scales. While only six scales were analysed quantitatively with the GelSight scanner, the absence of the AP type of mechanoreceptor from scales within the ventral and frontal planes suggests a difference in function (see below).

This study reveals that when the mechanoreceptor types are combined, *Hydrophis major* has approximately 7,000 scale mechanoreceptors distributed over the entire head. The texas rat snake, *Elaphe obsoleta lindheimeri*, is the only other snake where the total number of cephalic skin mechanoreceptors has been reported and has approximately 6,000 receptors over its head (Jackson & Sharawy, 1980). The similar number of mechanoreceptors over the head raises the question of what the relative importance of mechanoreception in comparison to other sensory modalities in snakes is. The star-nosed mole, *Condylura cristata*, for example, which relies entirely on mechanoreception to optimise foraging in its underground habitat possesses 25,000 mechanoreceptors on its nose (Catania, 1999). The American alligator, *Alligator mississippiensis* and the nile crocodile, *Crocodylus niloticus,* are both known for their superbly sensitive mechanosensory systems with 4,200 ± 94 and 2,900 ± 134 mechanoreceptors over the head, respectively (Leitch & Catania, 2012), but also for their visual acuity when foraging (Nagloo et al., 2016). Both the texas rat snake and *H. major* are situated between the numbers of the star-nosed mole and the crocodilians, which is striking considering their much smaller head size compared to crocodilians and may also be indicative of the importance of mechanoreception as a sense in snakes. Further research on the neural network, e.g. the trigeminal nerve, behind cephalic mechanoreceptors is required to fully understand the importance of mechanoreception in *H. major* and other snakes.

We found marked sexual dimorphism for all mechanoreceptor traits of SD mechanoreceptors for all six scales examined in this study (Figure 5; Supplementary Figure 3). Males had taller mechanoreceptors on the lateral and dorsal sides of the head, while females had taller mechanoreceptors anteriorly on the rostrum. Females generally had more numerous mechanoreceptors on the dorsal and anterior sides of the heads than males. Head size significantly influences all mechanoreceptor traits (Table 3), but head size was not statistically different between the male and female specimens sampled in this study, so sexual dimorphism in mechanoreceptor number and height is unlikely to be due to allometry.

Although note that only the head of one male specimen was imaged in its entirety in this study, we expect to find sexual dimorphism in the abundance and distribution of mechanoreceptors over the entire head if more individuals are examined. Sexual dimorphism in mechanoreceptor size and distribution has been observed previously in at least one other sea snake lineage, the turtle-headed sea snake, *Emydocephalus annulatus*, where it is likely associated with tactile mating behaviours given the distribution of larger mechanoreceptors over the chin and anal scale that are in frequent contact during courtship (Crowe-Riddell et al., 2021). Freshwater *Helicops* snakes, *H. pastazae* and *H. angulatus*, also show sexual dimorphism in coverage, density and number of cephalic mechanoreceptors (García-Cobos et al., 2021). Sexual dimorphism in sensory traits may be related to sexual selection to find mates (e.g., longer tongue-tines in American copperheads; (Smith et al., 2008), tactile courtship (see below) and/or ecological differences among sexes. Future studies should aim to sample both sexes during ontogeny to identify shifts in mechanoreceptor traits at sexual maturity.

It is unclear whether cephalic scale mechanoreceptors, in addition to being hydrodynamic and tactile organs, are homologous to body rugosities and would thus change in size during the breeding period when the scales of some sea snakes become increasingly rugose (Avolio et al., 2006a, 2006b). In many sea snake species, males are generally more rugose (possess rough scales) along their body than females (Avolio et al., 2006a), and rugosity increases during the breeding season when these rough scales are hypothesised to confer a better grip on the female by the male (Avolio et al., 2006b). In some snakes, the importance of mechanoreceptors along the chin of males has been observed during mating, where the male repeatedly rubs his chin along the female’s back (Crowe-Riddell et al., 2021; Noble, 1937; Senter et al., 2014). Although we do not report particularly high mechanoreceptor values on the genial scale sampled for *H. major*. It is unclear whether the scales containing scale mechanoreceptors would undergo any change during the breeding period in sea snakes (Crowe-Riddell et al., 2021). The particularly high abundance and the differences in the size of the SD mechanoreceptor type in other *Hydrophis* sea snakes may also be due to ecological or sexual selection pressures, however, there is a dearth of studies on intraspecific variation in mechanoreceptor traits to test these hypotheses.

### 4.3. Variation in mechanoreceptor distribution over individual scales

When observing the arrangement of mechanoreceptors on individual scales, both SD and AP types often appear with the relative proportion of mechanoreceptor types varying in different scale types and/or the AP type entirely lacking (Figure 7). This may be due to each type providing different sensory information. Some species of snakes, geckos and iguanids show the same pattern as *H. major*, where for all scales except the rostrum, the scale mechanoreceptors occur at higher densities along the edges of the scales (Jackson & Sharawy, 1980; Matveyeva & Ananjeva, 1995; Riedel & Schwarzkopf, 2022). The lack of receptors along rostral scale edges may be due to their overlapping on their caudal edge (Ananjeva et al., 1991; Matveyeva & Ananjeva, 1995).

We predict that the distribution of scale mechanoreceptors in *H. major* follows an adaptive pattern to allow for an optimised sensitivity in the detection of either a hydrodynamic or mechanical signal. The alignment of scale mechanoreceptors into lines on many scales suggests a strong bias for signals emanating from specific directions. For example, the rostral SD mechanoreceptors may mask the signal of the caudal SD mechanoreceptors in the horizontal plane but may be strongly stimulated by signals perpendicular to this alignment, as would be expected for the high density of mechanoreceptors on the ventral end of scales lining the mouth. Similarly, the lateral scales of *H. major* show a dorso-ventral alignment of the mechanoreceptors along the scale edge, while mechanoreceptors on ventral and dorsal scales are aligned along the lateral edge of each scale, indicating different axes of sensitivity (Figure 4 and 7).

A striking example of this type of adaptive mechanoreceptor orientation could be the supralabial scales in *H. major*, which show a row of scale mechanoreceptors on the ventral edge of the scale, along the edge of the mouth (Figure 6), an arrangement that has also been observed in crocodilians (Leitch & Catania, 2012) and in some geckoes (Riedel & Schwarzkopf, 2022). All supra-and sublabial scales have the highest mechanoreceptor densities along the edge of the mouth (Figure 7). This distribution pattern may be facilitating tactile perception, as mechanoreceptors detect the orientation of a captured prey item, thereby enabling the animal to subsequently rotate the prey to be swallowed tail or head first (Udyawer et al., 2020; Voris & Voris, 1983). *Hydrophis major* has been recorded capturing a striped catfish (Berthomier, 2015; Guémas, 2020) that was then rotated within the snake’s mouth prior to being swallowed whole. In cobras, *Ophiophagus hannah*, and coral snakes, *Micrurus fulvius*, it was also found that the animals are able to detect the scale overlap of their prey items and thus orient the prey accordingly to swallow them head first (Greene, 1976). A similar process could be required for sea snakes targeting fish with scales or spines such as pufferfish.

### 4.4. Functional implications

As previously stated, we suspect that mechanoreceptors in *H. major* are able to detect hydrodynamic stimuli (Crowe-Riddell et al., 2016; Crowe-Riddell, Williams, et al., 2019; Thewissen & Nummela, 2008; Westhoff et al., 2005). We have identified two mechanoreceptor types that both have underlying dermal papilla indicative of Meissner-like corpuscles used for detecting mechanical stimuli via deformation of the skin elevation (Alesci et al., 2024; Crowe-Riddell, Williams, et al., 2019; Jackson & Doetsch, 1977b). These SD and AP mechanoreceptor types are different in the outer shape and location of their dermal papillae in relation to the highest point of the mechanoreceptor above the surface of the dermis (Figure 2). These differences may indicate specialised functions for each of the mechanoreceptor types. SD mechanoreceptors all have the same orientation, while AP mechanoreceptors have their dermal papilla positioned rostrally and before the highest point of the mechanoreceptor, giving AP types a very specific orientation.

We hypothesise that AP types may serve as proprioceptors and be crucial in the positioning of the head in relation to the rest of the body, while SDs may be more suited to mediate mechanoreception and hydrodynamic perception. The different shapes of the mechanoreceptors most likely make them subject to different hydrodynamic stimuli and to different pressure regimes above the mechanoreceptor, similar to the pressure regime above the fin of a shark but on a much smaller scale (Xu et al., 2021).

One can assume that, similar to fish, the direction of the contact of a pressure signal in the water column with an AP will depend on the positioning of the head while swimming or in relation to pre-existing water currents and thus allow the snake to correct the position of its head within its environment (Porfiri et al., 2022). SD mechanoreceptors lack directionality and thus must have the potential to detect signals from any direction. However, it is important to note that, while AP mechanoreceptors, which possess directionality due to their asymmetrical morphology, and SD mechanoreceptors, which display a radial symmetry, may modify local hydrodynamic flow (i.e. boundary layer). Hydrodynamic pressure regimes may therefore be significantly influenced by the arrangement of the mechanoreceptors with respect to each other and specific sensory requirements. APs may play a role in striking prey, as this action requires precise positioning and movement of the head, should their role in proprioception be confirmed. However, studies on the hydrodynamics of snakes are rare and mainly focus on locomotory propulsion (Iosilevskii & Rashkovsky, 2020) and prey striking behaviour (Segall et al., 2019; Van Wassenbergh et al., 2009) and thus more biomechanical research needs to be conducted on whether the variation in distribution of cephalic mechanoreceptors affect the hydrodynamics of a sea snake and how detection of a signal in these mechanoreceptors occurs.

While it is unclear what the specialised functions of each of these two mechanoreceptor types provides, we suspect that both respond to hydrodynamic stimuli (Thewissen & Nummela, 2008; Westhoff et al., 2005). Indeed, even though *Hydrophis major* has relatively large mechanoreceptors, other species of sea snakes have much smaller mechanoreceptors (Crowe-Riddell, Williams, et al., 2019) which may indicate that the size and distribution of mechanoreceptors is influenced by each species’ ecology and behaviour. For example, different mechanoreceptor traits might be linked to water turbidity/visibility or predatory hunting strategies. Tall AP mechanoreceptors might increase sensitivity for hunting schooling fish in open water, while higher numbers of SD mechanoreceptors might increase resolution for close-range differentiation while ambush hunting for crevice or burrow dwelling prey (e.g. Gobidae, Anguilliformes). It is expected that species of sea snakes that dive to deep waters, where little sunlight penetrates (Crowe-Riddell, D’Anastasi, et al., 2019; Speed et al., 2022), or live in highly turbid environments such as estuaries (Voris, 2015), may rely more heavily on hydrodynamic perception to “see” as the eyes would not be as effective in these conditions, in a similar way to the star-nosed mole (Catania, 1999). It is therefore reasonable to assume that the ecological behaviour of sea snakes has influenced the evolution of the size and distribution of their scale mechanoreceptors to meet their varying sensory needs.

## 5. CONCLUSION

Our study has identified two distinct types of scale mechanoreceptors in a sea snake, one smooth and rounded dome-shaped mechanoreceptor found in other snakes, the other is larger and forms an asymmetrical peak, which has not been described previously. They can easily be distinguished by their histology, outer shape, and height. Analyses of these morphological characteristics have enabled a better understanding of the complexity of cephalic mechanoreceptor distribution and allowed us to make predictions on structure-function relationships in snakes, specifically that these two mechanoreceptor types are functionally different in their sensitivity to hydrodynamic stimuli. Our study shows that gel-based stereo profilometry is a rapid and accurate tool to quantify and measure cutaneous neurobiological structures. It has allowed the creation of the first distributional map of mechanoreceptor density in sea snakes. The clear patterns in distribution, with frontal and labial scales showing the highest densities of mechanoreceptors, as well as the increased size of their receptors, may indicate complex ecological adaptation to marine environments.

## Supporting information

All supplementary material

## 6. ETHICS

Snakes were collected and euthanized under approval from The University of Adelaide Animal Ethics Committee (Science) (Approval number S/2021-017) or measurements were taken from deceased animals that were alcohol-preserved at the South Australian Museum.

## 7. AUTHOR’S CONTRIBUTIONS

AW, JMC-R and SPC conceived of the study. AW conducted dissections and gel-based profilometry with assistance from JMC-R. AW carried out microscopy analyses with assistance from CJ and MHH. CJ wrote code for heatmap of mechanoreceptors over the head. AW conducted data analyses with input from JMC-R, KLS and SPC. JMC-R and KLS collected sea snake specimens. AW and JMC-R wrote the manuscript with input from all co-authors.

## ACKNOWLEDGEMENTS

For access to museum specimens, we thank Ralph Foster from the South Australian Museum. We would like to thank Chris Leigh, Jane Sibbons (Adelaide Microscopy, South Australia), Julian Ratcliffe (Bioimaging Platform, La Trobe University) for assistance and training provided for the ultramicrotome, slide scanner and scanning electron microscope. We are grateful to Caroline Kerr (La Trobe University) for supporting AW during her time at La Trobe University. We would like to thank Vohn Garcia for allowing us to use one of his *Hydrophis major* images. We acknowledge Koundinya Desiraju for the use of part of his R script from github (https://github.com/koundy/ggplot_theme_Publication).

