## Supplementary material for "Morphology and distribution of a new scale mechanoreceptor type in olive-headed sea snakes (*Hydrophis major*)": All supplementary material

Supplementary table 1: Thresholds set to each cephalic scale sample in order to distinguish asymmetrical peak and smooth dome mechanoreceptor types. Only when both the height range threshold and the roundness threshold match in a mechanoreceptor can it be associated to either a smooth dome or asymmetrical peak type. The height range threshold corresponds to the fraction of the height range of the receptors on a given scale above which mechanoreceptors are asymmetrical peak type. The roundness threshold is the threshold beneath which receptors of a given scale are of the asymmetrical peak type. Any particle outside of these constraints was considered a smooth dome type mechanoreceptor.

| Cephalic scale | Height range threshold | Roundness threshold |
|---|---|---|
| Genial scale | 0.92 | 0.6448 |
| Postocular scale | 0.81 | 0.8687 |
| Nasal scale | 0.92 | 0.8687 |
| Supralabial scale | 0.92 | 0.8687 |
| Rostrum | 0.92 | 0.5915 |
| Parietal scale | 0.92 | 0.8687 |

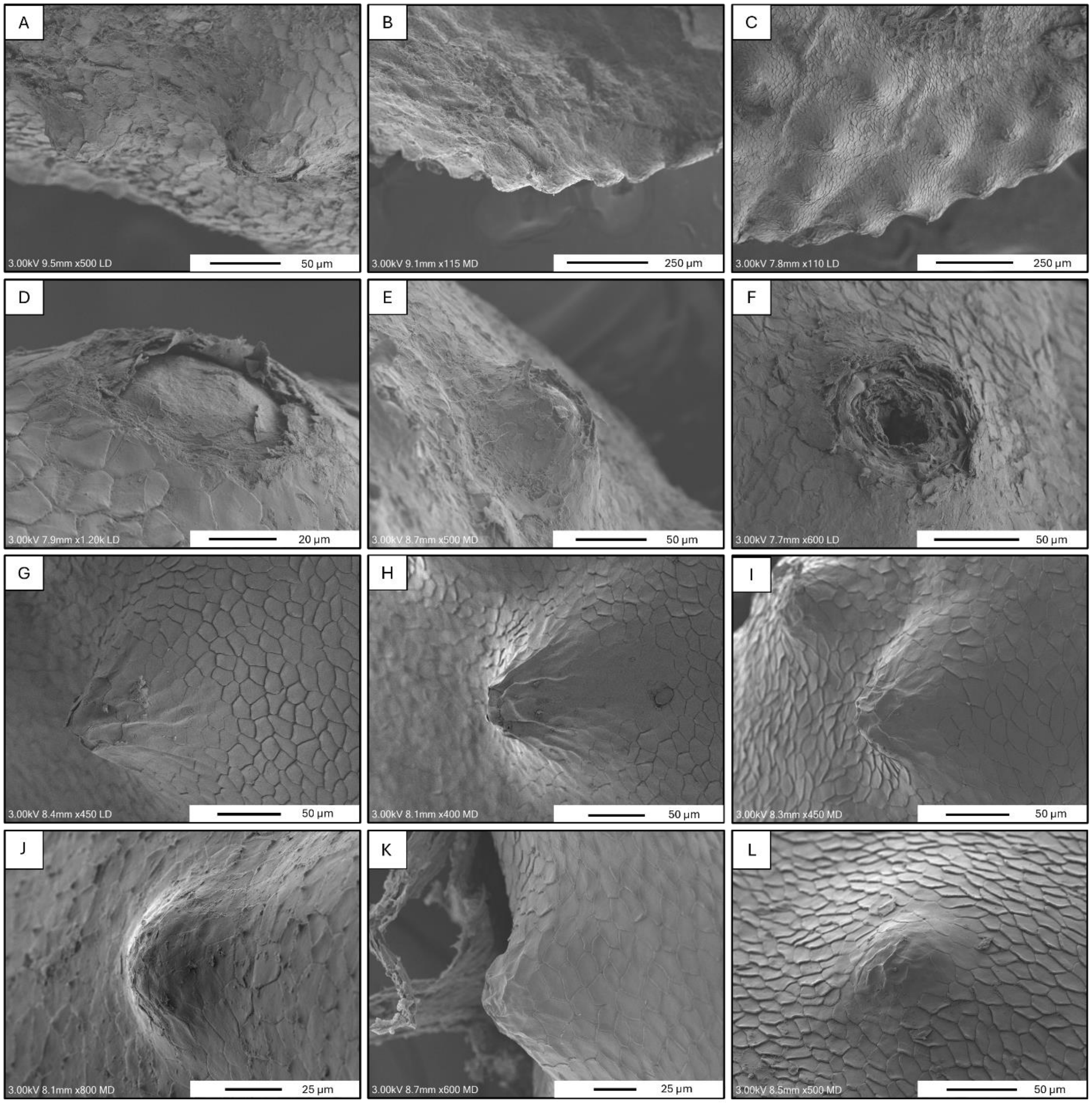


<u>Supplementary figure 1:</u> Scanning electron microscope images of scale mechanoreceptors across different scales in *Hydrophis major*. (A) was obtained on a supralabial scale and shows size differences between scale mechanoreceptors. (B) and (C) show the alignment of scale mechanoreceptors along the edge of the supralabial scale. (D), (E) and (F) show various stages of damage to the scale mechanoreceptors and were obtained on the parietal and nasal scales. (G), (H) and (I) are asymmetrical peak mechanoreceptors on the postocular scale. (J), (K) and (L) are smooth domes obtained on the parietal and postocular scales. The image legend in the lower left corner gives the voltage, the working distance, the magnification and an abbreviation for each detector used. A scale bar for each image is given in the lower right corner.

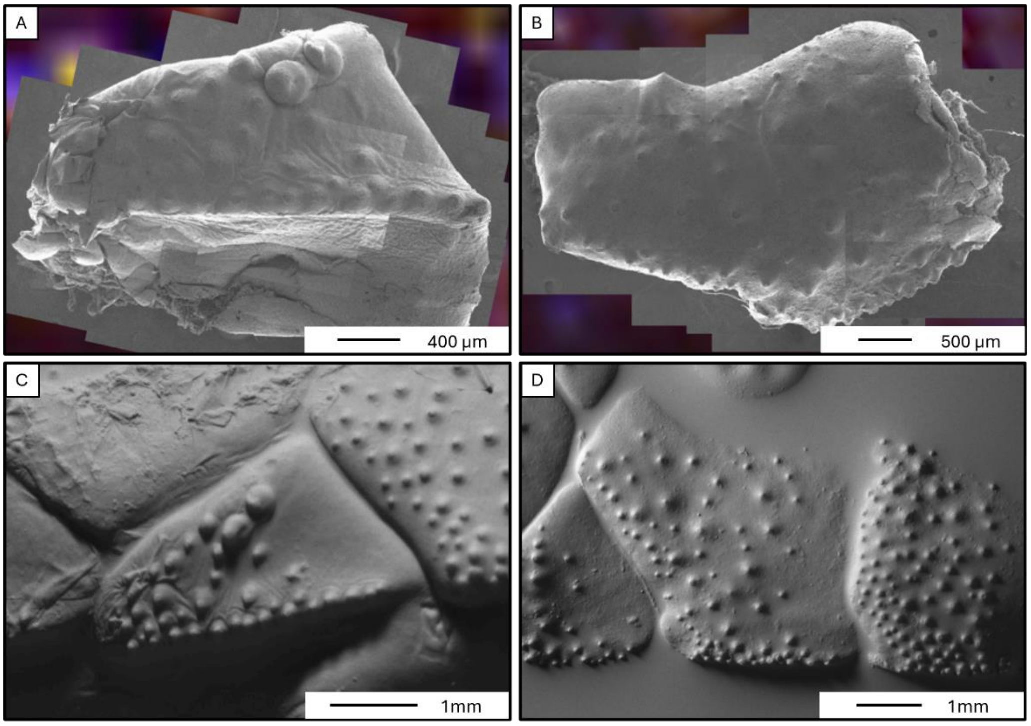


Supplementary figure 2: Side-by-side comparison of the same two supralabial scales imaged using scanning electron microscopy (A, B) and the GelSight scanner (C, D).

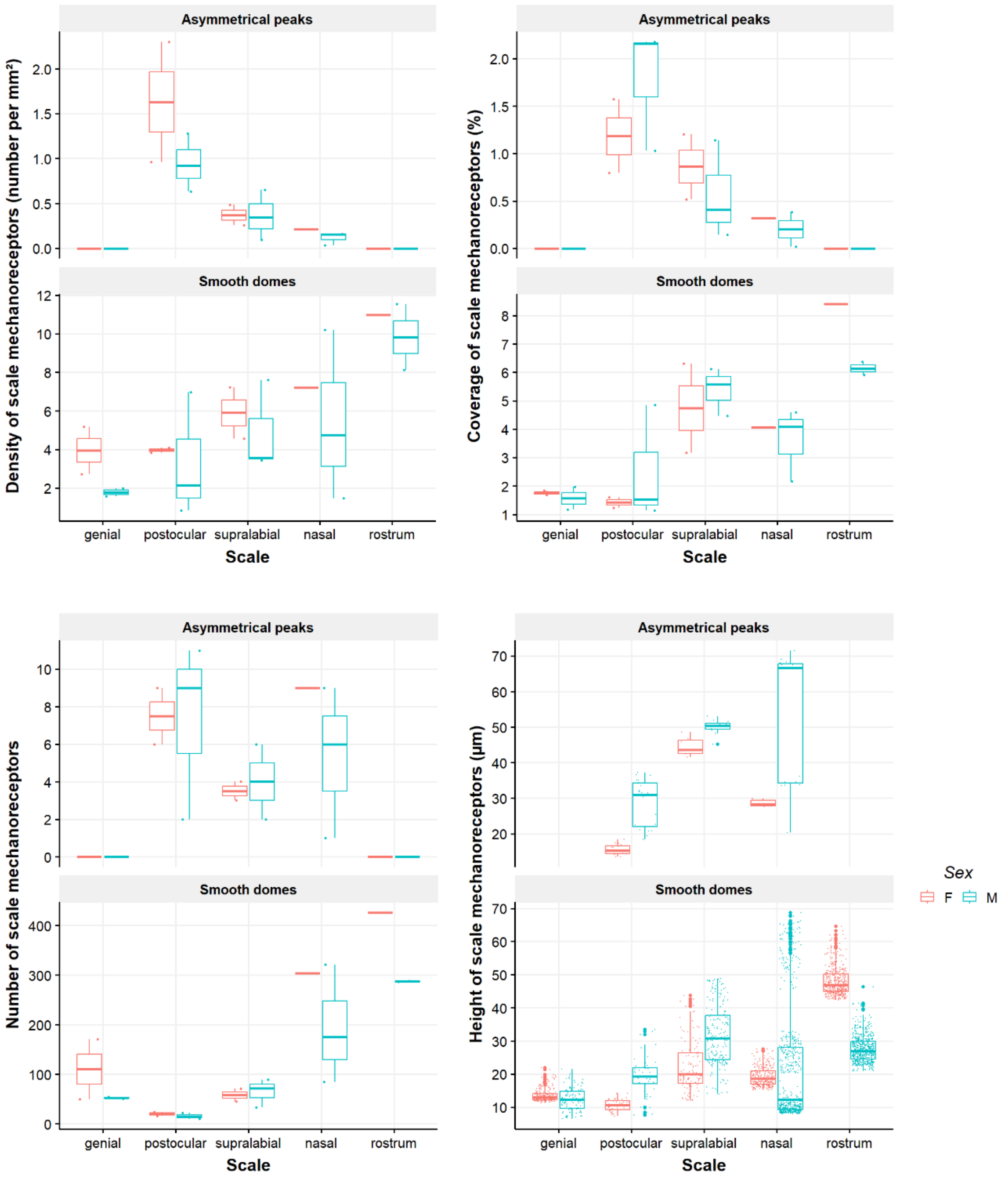


Supplementary figure 3: Boxplot of density (per mm²), coverage (%), number and height (µm) of the two types of mechanoreceptor per scale differentiated by sex in *Hydrophis major* across five cephalic scales. The two types of mechanoreceptors are smooth domes and asymmetric peaks. The colours differentiate males (M) and females (F). Each box represents the interquartile range (IQR), with the line inside the box marking the median. Outliers are presented as large points. For the density of mechanoreceptors, the points represent the mechanoreceptor density per individual. For the coverage, the points represent the mechanoreceptor coverage per individual. For the number of mechanoreceptors, the points represent the total number of receptors per individual, note the different y axis. For the height of mechanoreceptors, the points represent individual receptor heights.

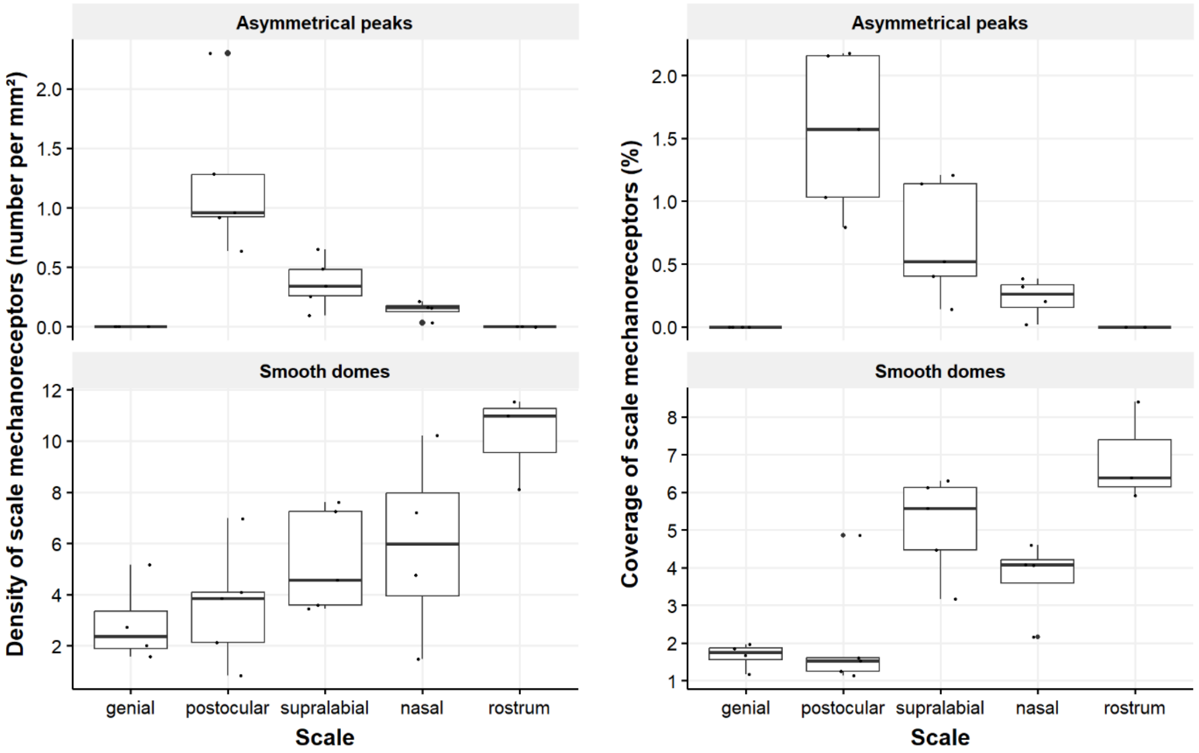


Supplementary figure 4: Boxplot of the density (per mm²) and the coverage (%) of the two types of mechanoreceptor (smooth dome and asymmetric peak) per scale in *Hydrophis major* across five cephalic scales. Each box represents the interquartile range (IQR), with the line inside the box marking the median. Outliers are presented as large points. Statistical analysis using the Kruskal-Wallis test revealed significant differences in the density of mechanoreceptors between scales (chi-squared = 2148, df = 4, p-value $< 2.2e^{-16}$). The points represent the density of mechanoreceptors per specimen per scale. For the coverage of mechanoreceptors, the points represent the coverage of mechanoreceptors per specimen per scale. Statistical analysis using the Kruskal-Wallis test revealed significant differences in the coverage of mechanoreceptors between scales (chi-squared = 3133.7, df = 4, p-value $< 2.2e^{-16}$).
